# New approach and new program for analyses of false negatives-contaminated data in medicine and biology

**DOI:** 10.1101/660324

**Authors:** Jaroslav Flegr, Petr Tureček

**Author notes:** Corresponding author: Department of Philosophy and History of Science, Faculty of Science, Charles University, Viničná 7, Prague, 128 43, Czech Republic.

## Abstract

**Background:** No serological assay has 100% sensitivity. Statistically, the concentration of specific antibodies against antigens of parasites decreases with the duration of infection. This can result in false negative outputs of diagnostic tests for the subjects with old infectiong, e.g., for individuals infected in childhood. When a property of seronegative and seropositive subjects is compared under these circumstances, the statistical tests can detect no significant difference between these two groups of subjects, despite the fact that infected and noninfected subjects differ. When the effect of the infection has a cumulative character and subjects with an older infection (potential false negatives) are affected to a greater degree, we can even get paradoxical result of the comparison – the seronegative subjects have on average lower value of certain traits, e.g. IQ, despite the infection having a negative effect on the trait. A permutation test for the contaminated data, implemented, e.g., in the program Treept or available as a comprehensibly commented R function in the supplement of this paper, can be used to reveal and to eliminate the effect of false negatives.

**Methods:** We used a Monte Carlo simulation in the program R to show that the permutation test implemented in the programs Treept and PTPT is a conservative test.

**Results:** We showed that the test could provide false negative but not false positive results if the studied population contains no subpopulation of false negative subjects. We also introduced R version of the test expanded by skewness analysis, which helps to estimate the proportion of false negative subjects based on the assumption of equal data skewness in groups of healthy and infected individuals.

**Conclusions:** Based on the results of simulations and our experience with empirical studies we recommend the usage of permutation test for contaminated data whenever seronegative and seropositive individuals are compared.

## Introduction

The reported decrease of specific antibodies with time from the onset of infection increases the risk of false negative test results in subjects with old infections, e.g., in individuals infected in childhood^1–3^. This is also true for parasites that stay dormant in infected cells until the end of the life of infected hosts. Any subsample of seronegative subjects could therefore be contaminated with an unknown proportion of misdiagnosed parasite-positive individuals who got infected a long time ago^4–6^. This subpopulation of infected but seronegative subjects could be the most influenced by the infection (Figure 1B) because of the long duration of their infection or because their infection took place in early stages of their ontogenesis. This could result in a paradox (Figure 1C). The seropositive subjects could have on average higher IQ scores (or higher body weight) while the intelligence (or body weight) of seropositive subjects declines with the assessed length of infection (obtained from clinical records or assessed by the level of antibodies).

**Figure 1.**
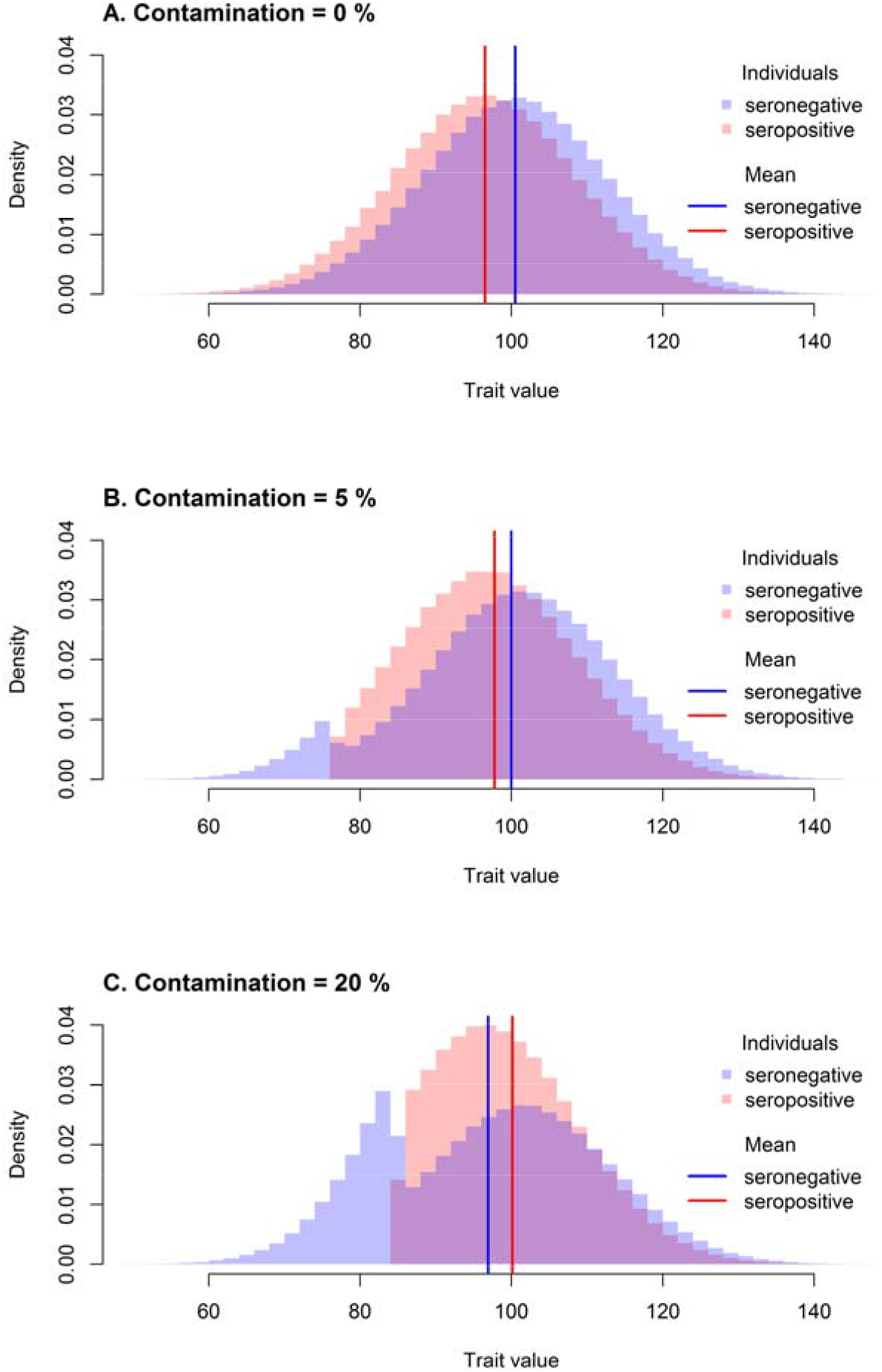
Exemplar distributions under 3 different contamination levels. The proportion of seropositive individuals (50%), the difference between healthy and infected individuals (5) and the standard deviation (10, corresponding to Cohen’s d = 0.5 in non-contaminated sample) within healthy individuals are held constant. Histogram C serves as a demonstration of the paradoxical result caused by a high contamination when the seronegative is a lower seropositive mean trait value despite the fact that healthy individuals score higher than infected individuals.

The contamination of a parasite-free subsample with false negative individuals can be revealed and eliminated by permutation tests with the reassignment of suspect cases between subsamples^4, 5^. Such permutation tests can be performed using the program Treept, originally called PTPT^7, 8^ modified for an analysis of data contaminated with an unknown number of subjects with false negative diagnosis using the method of reassignment of potentially false negative subjects ^4^. This freeware program is available at http://web.natur.cuni.cz/flegr/treept.php. The updated version of the test suited for R can be found in the supplementary material of this paper in the form of comprehensibly commented R script.

The algorithm of the one-tailed permutation test with data reassignment is as follows: Particular percentage (e.g. 5, 10, 15, 20 or 25 %) of subjects with the lowest (highest) value of the dependent variable, for example IQ score, is relocated from the group of parasite-seronegative subjects to the group of the parasite-seropositive subjects. Then, the difference of means of these two groups is calculated. In the next 9,999 steps, the empirical values of the analysed variable are arbitrarily assigned into two groups held at the size of the original seronegative and seropositive groups. The particular percentage of cases with the lowest (or highest) values of the focal variable (e.g. IQ) in the pseudoseronegative group is relocated to the pseudoseropositive group, and the difference between the means of the two groups is calculated. Finally, all 10,000 differences (including the one calculated from non-permuted data) are sorted from highest to lowest. The percentage of the differences higher or equal to that calculated on the basis of the non-permuted data is considered as the statistical significance (p) – the probability of obtaining the same or higher difference between the means of two groups, if the null hypothesis is correct and subjects are assigned into seropositive and seronegative groups randomly.

Our main aim is to show that the permutation test for contaminated data does not provide false positive results, i.e., it does not return lower p than a standard permutation test if no false negative subjects exist in the studied population. The second aim is to develop a new tool for the skewness analysis, which can be used to estimate the approximate proportion of false-negative subjects in the studied population.

## Methods and Results

A Monte Carlo simulation was performed with R 3.3.3. We generated a population of 150 parasite-free and 150 infected subjects (mean intelligence was 101.5 in the parasite-free group and 98.5 in the infected group – the between-group difference was 3, the population mean intelligence was 100). Subjects were normally distributed around group means with equal standard deviations (SD). We used different SDs (6, 9, 12, 15, 30) corresponding to different effect sizes expressed by Cohen’s d (0.5, 0.33, 0.25, 0.2, 0.1). Then we ran a standard permutation test. We randomly permutated the infection status of all subjects 10,000 times and calculated a fraction of permutations where the difference between two groups (pseudo-parasite-free and pseudo-parasite-infected subjects) was equal to or larger than the difference between the groups in non-permutated data (p value of a standard permutation test). Then, we repeated the analysis using a one-tailed permutation test for contaminated data. Namely, after the generation of sets of parasite-free and parasite-infected subjects (or after the generation of sets of pseudo-parasite-free and pseudo-parasite-infected subjects by permutation of the infection status), we relocated 5, 10, 15, 20, 25, 30 or 50% of subjects with the lowest intelligence from the parasite-free (or pseudo-parasite-free) set to the parasite-infected (or pseudo-parasite-infected) set. Again, we calculated a fraction of permutations with the difference between the groups equal to or larger than the value computed for the non-permuted data (p values of the permutation test for contaminated data). We used populations generated for the standard permutation test (each initial population was used once for each fraction of relocated subjects). In total, 10,000 original populations were generated for each SD, therefore 10,000 independent permutation tests were conducted for each combination of SD and each relocated fraction. The resulting p values were averaged over permutation tests with the same population SD and the same relocated fraction. The results are shown in the Table 1. With the proportion of relocated subjects, the average p-value grew for every standard deviation. The visualization of this growth can be found in Figure 2A. In this figure, the p-value of the standard permutation test was subtracted from each p-value of the permutation test for contaminated data (negative values therefore correspond to a decrease, and positive to an increase, of p-value in comparison to a standard permutation test).

**Table 1.**
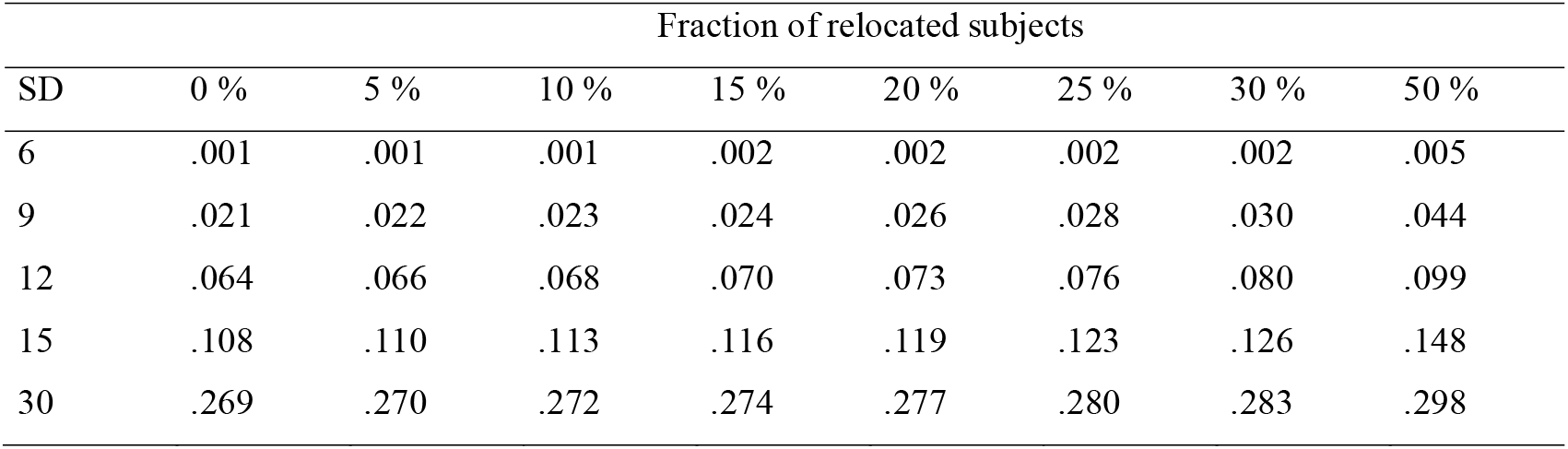
Effect of relocation of hypothesized false negative subjects on the results of a permutation test if no such subjects exist in the population. The table shows p-values computed with the permutation test for contaminated data when the population under study contains no false negative subjects. The simulation experiments were performed on populations that differ by variances (rows) with the relocation of different fractions of IQ-lowest individuals (columns) from the high-IQ (seronegative) group to the low-IQ (seropositive) group. The first column (0%) shows the (most significant) results of permutation tests performed without any relocation of data. For details see the Methods section. The fixed effect was 3 IQ points. The population size was 300, and the proportion of seropositive individuals in the original sample (0% relocation) was 0.5.

| SD | Fraction of relocated subjects |  |  |  |  |  |  |  |
| --- | --- | --- | --- | --- | --- | --- | --- | --- |
|  | 0 % | 5 % | 10 % | 15 % | 20 % | 25 % | 30 % | 50 % |
| 6 | .001 | .001 | .001 | .002 | .002 | .002 | .002 | .005 |
| 9 | .021 | .022 | .023 | .024 | .026 | .028 | .030 | .044 |
| 12 | .064 | .066 | .068 | .070 | .073 | .076 | .080 | .099 |
| 15 | .108 | .110 | .113 | .116 | .119 | .123 | .126 | .148 |
| 30 | .269 | .270 | .272 | .274 | .277 | .280 | .283 | .298 |

**Figure 2:**
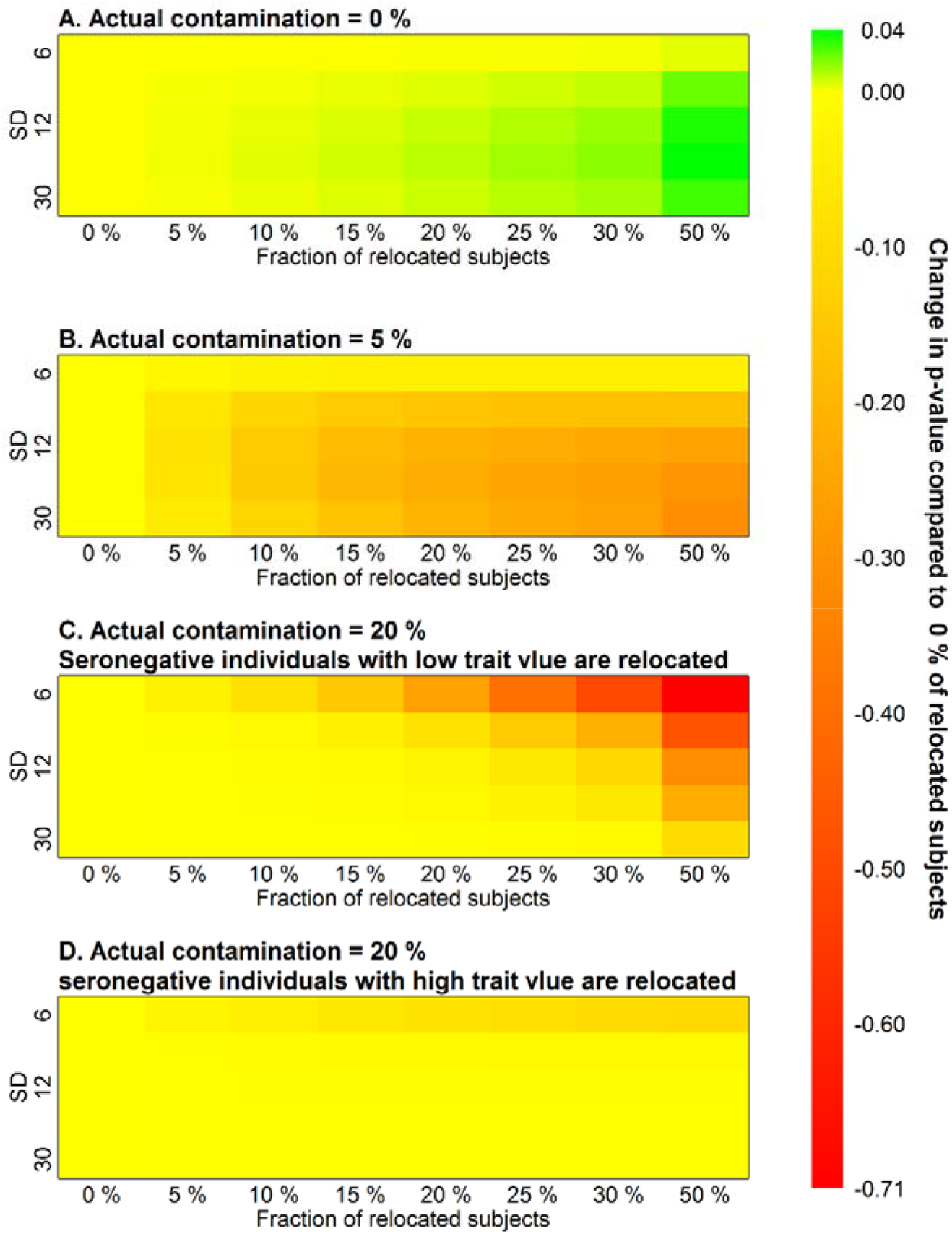
Heatmap of the average difference between the p-value of standard permutation test and p-value of the respective permutation test for contaminated data. One-tailed tests were used. The p-value increases with the fraction of relocated individuals if no actual false negative individuals are present (A) and decreases if the sample is contaminated (B, C). This is true even if the wrong relocation direction is employed due to a paradoxical switch in the order of group means (D).

When several exceptional data points (outliers) are present, the p-value of one or more contamination levels can be lower than p-value for 0% contamination. This is more frequent when the effect size is very small and the p-value fluctuates due to a larger impact of random noise in the data. The probability of a p-value being higher for a certain proportion of relocated data than the p-value of a standard permutation test in a particular simulation run was evaluated for each level of contamination and SD from the set of generated data described above. The results are reported in the Table 3 and shown in the Figure 3A.

**Figure 3:**
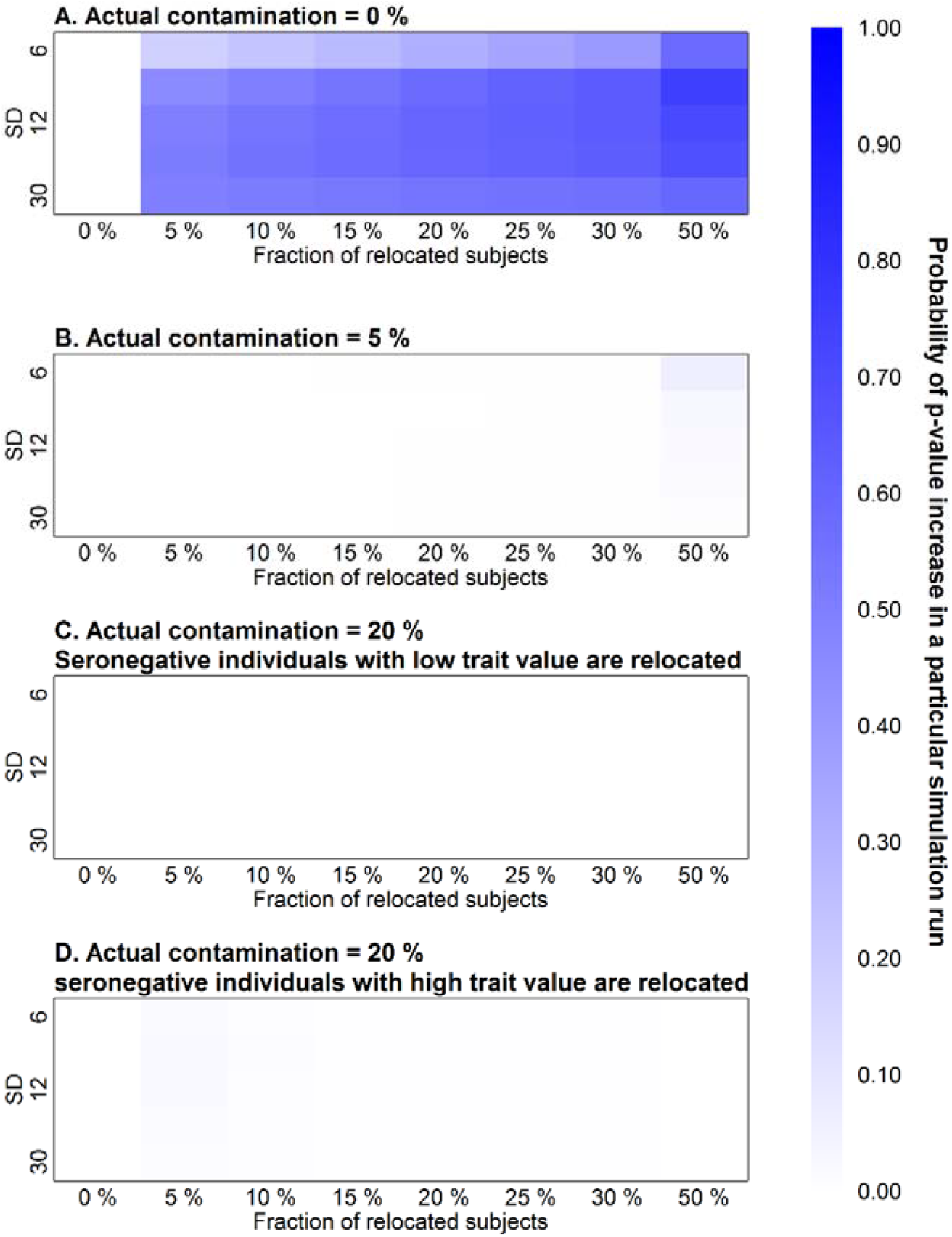
Heatmap of probability of p-value increase in particular simulation run. The p-value does not increase in 100% of non-contaminated samples when a permutation test for contaminated data is used, but the probability that it happens is very high compared to samples in which false negative individuals occur.

For comparison, the same computer simulation was conducted for a population of 150 seropositive and 150 seronegative individuals where 5% of seronegative individuals were false negative individuals with extremely low intelligence (example in Figure 1B). The average p-values of the permutation test for contaminated data are in Table 2. The graphical representation of the difference between a p-value for 0 % of relocated subjects and other contamination levels is represented in Figure 2B, and the probability of increase of p-value is shown in Table 4 and Figure 3B. In this case, the p-value decreases with the proportion of relocated individuals as expected.

**Table 2.**
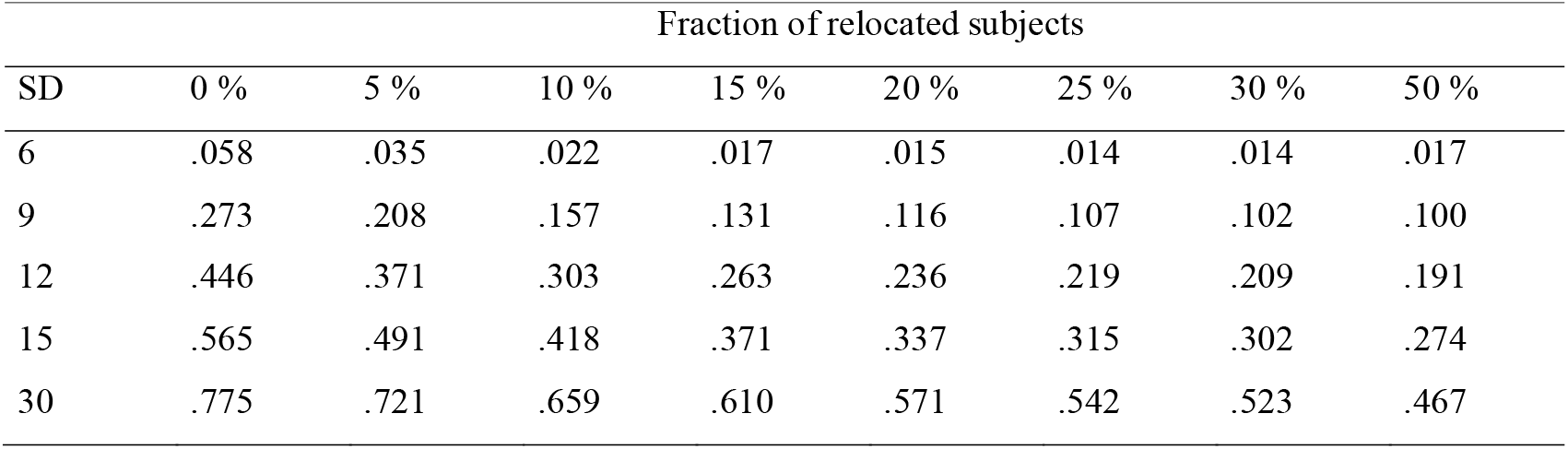
Effect of the relocation of hypothesized false negative subjects on the results of a permutation test if 5% of such individuals is present in seronegative group The table shows p-values computed with the permutation test for contaminated data when the seronegative group contains 5% of false negative subjects. The simulation experiments were performed on populations that differ by variances (rows) with relocation of different fractions of IQ-lowest individuals (columns) from the high-IQ (seronegative) group to the low-IQ (seropositive) group. The first column (0%) shows the (least significant) results of permutation tests performed without any relocation of data. For details see the Methods section. The fixed effect was 3 IQ points. The population size was 300, and the proportion of seropositive individuals in the original sample (0% relocation) was 0.5.

**Table 3.** The probability of a p-value being higher in a particular simulation run than a p-value with 0 % of relocated individuals. No false negative subjects are present in the population. The probability that p-value will increase for specified fraction of relocated individuals in a particular simulation run as compared to 0 % of relocated seronegative individuals. The simulated population are identical to the population represented in Table 1. The graphical summary can be found in Figure 3A. The relatively small numbers in first row are caused by the fact that in many simulation runs p-values remained <.0001 for a small fraction of relocated subjects when the effect size is relatively large.

| SD | Fraction of relocated subjects |  |  |  |  |  |  |
| --- | --- | --- | --- | --- | --- | --- | --- |
|  | 5 % | 10 % | 15 % | 20 % | 25 % | 30 % | 50 % |
| 6 | .18 | .23 | .27 | .31 | .35 | .39 | .58 |
| 9 | .45 | .50 | .54 | .58 | .61 | .64 | .75 |
| 12 | .50 | .54 | .57 | .60 | .62 | .64 | .71 |
| 15 | .51 | .55 | .57 | .59 | .61 | .63 | .68 |
| 30 | .50 | .52 | .53 | .54 | .55 | .56 | .59 |

**Table 4.** The probability of p-value being higher in particular simulation run than p-value with 0 % of relocated individuals. 5% of seronegative group are false negative individuals. The probability that a p-value will increase for a specified fraction of relocated individuals in a particular simulation run as compared to 0 % of relocated seronegative individuals. The simulated population are identical to those represented in Table 2. The graphical summary can be found in Figure 3B.

| SD | Fraction of relocated subjects |  |  |  |  |  |  |
| --- | --- | --- | --- | --- | --- | --- | --- |
|  | 5 % | 10 % | 15 % | 20 % | 25 % | 30 % | 50 % |
| 6 | .00 | .00 | .00 | .00 | .00 | .00 | .06 |
| 9 | .00 | .00 | .00 | .00 | .00 | .00 | .03 |
| 12 | .00 | .00 | .00 | .00 | .00 | .00 | .02 |
| 15 | .00 | .00 | .00 | .00 | .00 | .00 | .02 |
| 30 | .00 | .00 | .00 | .00 | .00 | .00 | .01 |

Two equivalent simulations were run to demonstrate the permutation test for contaminated data on the paradoxical dataset with a high proportion of false negative individuals. The first population of 150 seropositive and 150 seronegative individuals where 20% of seronegative subjects were false negative individuals with extremely low intelligence (Figure 1C). A similar one-tailed permutation test as in previous simulations was run as it was hypothesised that the average trait value of the healthy group is actually higher despite the paradoxical situation. The graphical representations of the results are in Figure 2C and Figure 3C. The second test with the same sample generation algorithm (150 seropositive, 150 seronegative, 20% false negative) was set to follow the default setting of the permutation test for contaminated data, which assumes the non-paradoxical situation and therefore relocates seronegative individuals with high trait value, thus widening the gap between the groups. Yielded p-value of one-tailed permutation test is then the proportion of random samples after relocation where the difference between groups (seronegative - seropositive) was lower than in the original sample (Figure 2D and Figure 3D). In both simulation tests on a sample with 20% contamination, the p-value of respective one-tailed permutation test decreased, so this sample was clearly distinguishable from the case in which no false negative subjects were present.

The appropriate direction of subject relocation can be determined on the basis of a skewness analysis of the original sample, which is available in the R version of the test^9^ if a parameter skewness.analysis is set to TRUE. The skewness analysis and its usage for the assessment of group mean order as well as the contamination level estimation is described in the Appendix of this paper. Using a two-tailed test is also worth consideration in this case. The p-value is then declining with the proportion of relocated individuals in all cases where false negative individuals are present (in well identified paradoxical situations only after the group means change their order into the right direction).

## Conclusions

The results of simulation showed that the permutation test for contaminated data does not provide more significant results than a standard permutation test if the experimental data does not contain a subpopulation of false negative subjects. This test is conservative when its usage is not necessary and allows one to avoid false negative results in the case of data contamination. This is due to the higher difference between the relocated seronegative and the original seropositive group in the presence of false negative data. The referential set of permutations with relocation remains the same in both cases, while the relative change in inter-group difference after relocation maintains an intermediate position between those two options (see Figure 4). Therefore, the positive result of this test, i.e. lowering the p-value with the growth of the proportion of relocated individuals, itself supports the hypothesis that the set of seemingly parasite-free subjects contains false negative subjects, who, most probably, have become infected a long time ago or in very young age.

**Figure 4.**
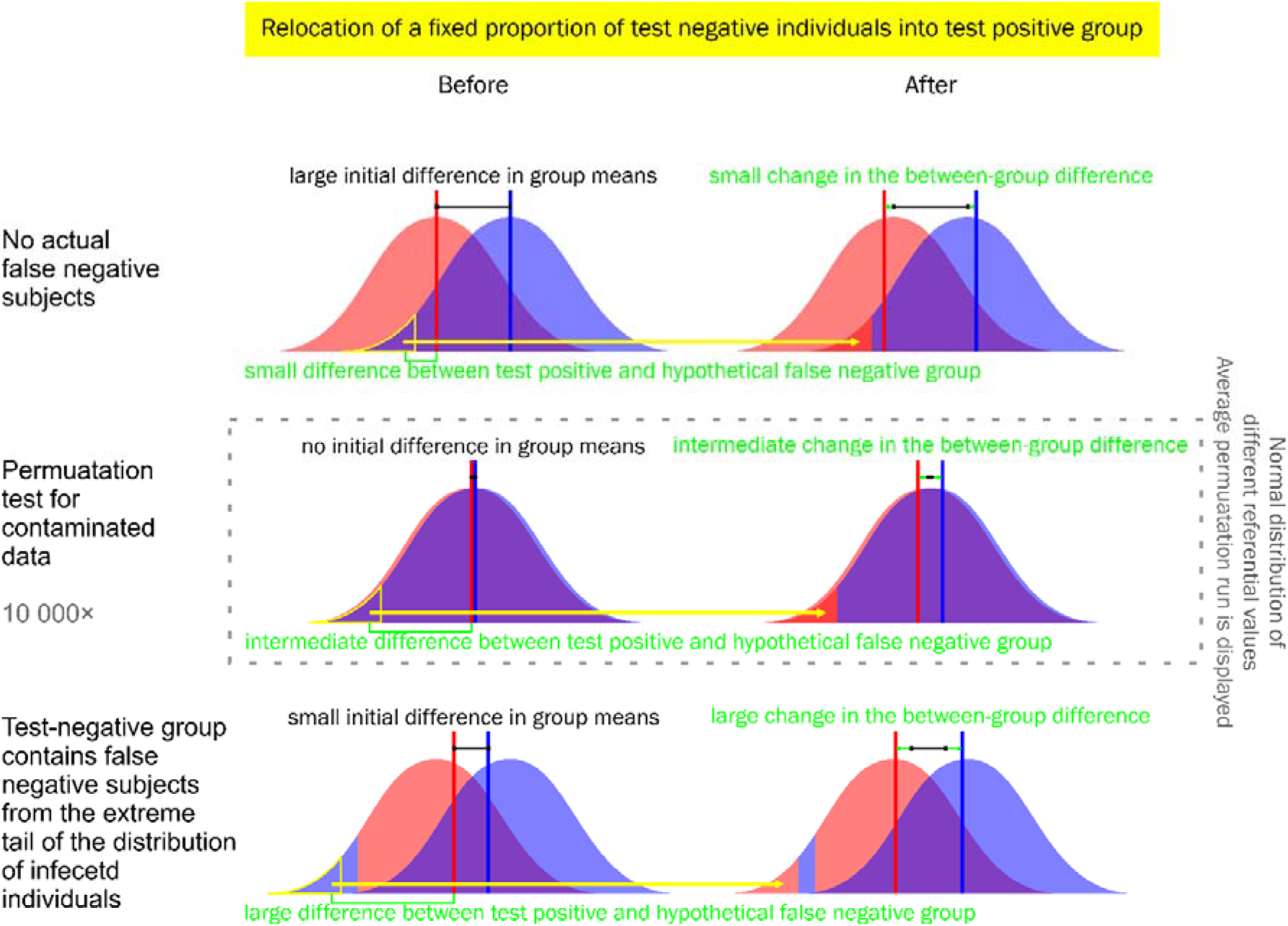
Graphical demonstration of the intermediate position of referential permutations with relocation between empirical cases of relocation of seronegative healthy subjects and false negative infected subjects. The increase in p-value in the case of non-contaminated data is much smaller than the increase caused by possible contamination, which can completely wipe out the actual inter-group difference or even cause a paradoxical switch of the group mean order. (See Table 5 or the position of 0 in the legend of Figure 2.)

**Table 5.** Risk associated with different combinations of data and used permutation tests The pressure to avoid a high risk associated with regular permutation test will lead us to the universal utilization of permutation test for contaminated data whenever properties of seropositive and seronegative subjects are compared. When we conduct a skewness analysis for contaminated data (see the Appendix) prior to the permutation test, we can lower the risks further by justification of regular permutation test or informed setting of relocated fractions of seronegative individuals in permutation test for contaminated data.

|  |  | Test |  |
| --- | --- | --- | --- |
|  |  | regular permutation test | permutation test for contaminated data |
| Data | non-contaminated | No risk | Small risk of false negative results |
|  | contaminated | **High risk of false negative or paradoxical results** | No risk |

Based on the theoretical grounding described above and our experience with the research of two unrelated species of parasites, *Toxoplasma gondii* ^5, 10^ and human cytomegalovirus^6^, we strongly recommend the usage of permutation tests for contaminated data^9^ whenever any properties of parasite-infected and parasite-free individuals are compared.

## Acknowledgements

We would like to thank Charlie Lotterman for his help with the final version of the paper.

## Appendix 1 Estimation of the fraction of false negatives by skewness analysis

The estimation of the actual contamination level is very difficult to discern and should be investigated more in future research. For now, we can seek assistance in a skewness analysis, which compares the skewness of trait value distribution in seropositive and seronegative groups. Skewness is defined as third standardized moment measuring the asymmetry of the probability distribution. We assume that healthy and infected individuals have an equally skewed trait distribution. This assumption is violated if false negative individuals are recruited from one of extreme tails of the distribution of infected individuals. If, for example, infected individuals with the lowest trait value are identified as seronegative (as seen in Figure 1B), the skewness of seropositive individuals becomes more positive and the skewness of seronegative individuals more negative. The exact opposite is true if individuals from the upper tail of the distribution are misdiagnosed as negative. The skewness comparison (available as a function in supplementary R script) of contaminated data compares Fisher-Pearson coefficient of skewness of seropositive and seronegative groups under different hypothesised contamination levels and returns the skewness values for each fraction of relocated subjects, p-values of the difference between them based on permutation test, the interval where the group skewness is not significantly different and a proportion of relocated seronegative individuals at which the difference between group skewness was smallest (i.e. the one the generated the most similarly skewed groups). This value generally underestimates the actual contamination, but any amendments would require additional assumptions about the distribution of healthy/infected individuals, which would not be necessarily met in empirical data. **Now we recommend the conduction a skewness comparison prior to the evaluation of the between-group difference and then the conduction of a permutation test for contaminated data for contamination levels between 0 and upper border of the interval, where the difference between group skewness was not significant.** We observed that the actual level of contamination in simulated data, where we can control the contamination level, rather closely matches the upper level of the similar-skewness interval due to the fact that the distributions of healthy and infected subjects largely overlap, and the extreme tail of seronegative distribution contains also extreme healthy individuals which are relocated prior to actual false negative individuals. For the same reason, however, we can suggest that the between-group difference for the relocated fraction where the group skewness are most similar (described above) closely matches the actual between-group difference in non-contaminated populations without false negative subjects.

The difference between skewness coefficients in seropositive and seronegative groups in the original sample without relocated individuals can also be evaluated in the R version of the permutation test for contaminated data^9^ (set skewness.analysis to TRUE). This analysis allows one to appropriately assess whether the seronegative group includes false negative subjects from the extreme tail of the distribution of infected individuals. By default, the permutation test for contaminated data assumes that the observed order of mean values of seropositive and seronegative groups accurately reflects the state of things in correctly determined healthy and infected groups. Therefore, the function will gradually relocate individuals from the lower tail of the distribution if the seronegative mean trait value is higher than the seropositive mean and vice versa (this can be changed by the parameter higher.healthy). If we do not alter default setting in paradoxical situations (Figure 1C), in which the order of group means was changed due to contamination, the test algorithm will increase the difference between the groups by relocating healthy individuals from the upper tail of distribution of seronegative subjects. The p-value will most likely decrease with the growing fraction of relocated individuals, as in other cases where false negative individuals are present. This might lead to a radical misinterpretation of the data (confirmation of the assumption of higher trait value in group of infected individuals) if attention is not payed to the skewness analysis. The skewness analysis of the original sample is not fooled as easily since mismatching the extreme tail of infected individuals as seronegative will alter the skewness of both groups substantially. Under an extreme proportion of false negatives, the skewness of both groups might actually be shifted in the same direction (e.g. positive in cases similar to the example in Figure 1C). However, the skewness of seropositive group will be still substantially more deflected than the skewness of the seronegative group, so the skewness analysis will return reliable results.

## Supplements (R scripts)

(Jaroslav Flegr and Petr Tureček, New approach and new program for analyses of false negatives-contaminated data in medicine and biology)

### Supplement 1

#### Permutation test for contaminated data

~~~
#####################################################
####hit ctrl+a and ctrl+r to install the function####
#####################################################
#This script contains contamination_perm_test() function
#contamination_perm_test(trait,identification,percentages=c(0,5,10),higher.
healthy=(mean(trait[identification==F])>mean(trait[identification==T])),run
s=10000,skewness.analysis=F)
#Arguments are described below.
#It is necessary to define funtion calculating skewness index first:
##Fuction that returns Fisher-Pearson coefficient of skewness.
##Input is a vector of numerical values.
FPskewness<-function(x){
return((sum((x-mean(x))^3)/length(x))/((sqrt(sum((x-
mean(x))^2)/length(x)))^3))
}
###contamination_perm_test
###Function that delegates the parameters to either one-tailed or two-
tailed tests described below
contamination_perm_test<-
function(trait,identification,percentages=c(0,5,10),higher.healthy=(mean(tr
ait[identification==F])>mean(trait[identification==T])),runs=10000,two.tail
ed=F,skewness.analysis=F){
if(two.tailed==F){
contamination_perm_test_one(trait=trait,identification=identification,perce
ntages=percentages,higher.healthy=higher.healthy,runs=runs,skewness.analysi
s=skewness.analysis)
}else{
contamination_perm_test_two(trait=trait,identification=identification,perce
ntages=percentages,higher.healthy=higher.healthy,runs=runs,skewness.analysi
s=skewness.analysis)
}
}
###This Function works with following arguments:
###trait - Numerical vector of trait values
###identification - Logical vector of assumed presence (T) or absence (F)
of infection
###percentages - Numerical vector of percentages of false negative amongst
negative subjects (contamination levels) for which the permutation test for
contaminated data will be run.
###two.tailed - Specifies the verison of the test, two.tailed=F is the
default.
###higher healthy - Logical. Indicates whether we assume the healthy
individuals to show higher (T) or lower (F) trait values. When not
specified, the script assumes this relationship based of group means with
no hypothesised contamination.
######It allows us to use the difference between the groups (not in
absolute values) in permutation test. In this scenario the seropositive
group mean is substracted from seronegative group mean and the one-tailed
permutation test is conducted accordingly.
###runs - Number of resamplings used in permutation test
###One-tailed version of the test
contamination_perm_test_one<-
function(trait,identification,percentages=c(0,5,10),higher.healthy=(mean(tr
ait[identification==F])>mean(trait[identification==T])),runs=10000,skewness.
analysis=F){
if(length(trait)!=length(identification)){
stop(“The vectors of trait values and infection indication are of different
lengths.”)
}
higher<- (mean(trait[identification==F])>mean(trait[identification==T]))
set.higher<-higher.healthy
orig.means<-tapply(trait,identification,mean)
orig.means<-data.frame(orig.means)
names(orig.means)<-“Original mean values”
rownames(orig.means)[which(rownames(orig.means)==“FALSE”)]<-“Identified as
healthy”
rownames(orig.means)[which(rownames(orig.means)==“TRUE”)]<-“Identified as
infected”
higher.report<-ifelse(higher==T,
“In original sample, individuals identified as healthy showed higher
\naverage trait value.”,
“In original sample, individuals identified as infected showed higher
\naverage trait value.”
)
concord<-ifelse(higher==set.higher,“Consequently,”,“Despite that,”)
set.report1<-paste(concord,ifelse(set.higher==T,
“healthy individuals were hypothesised to have higher \naverage trait value
in a contamination-free sample. \n”,
“infected individuals were hypothesised to have higher \naverage trait
value in a contamination-free sample. \n”
))
set.report2<-paste(ifelse(set.higher==T,
“\nFor each contamination level respective proportion of seronegative
\nindividuals with lowest trait value was relabeled as seropositive \nin
original sample as well as in each permutation test run.”,
“\nFor each contamination level respective proportion of seronegative
\nindividuals with highest trait value was relabeled as seropositive \nin
original sample as well as in each permutation test run.”
))
trait<-c(trait[identification==F],trait[identification==T])
infected<-sort(identification)
count.healthy<-sum(!identification)
count.infected<-sum(identification)
Nperc<-length(percentages)
vector.ident<-list()
for(i in 1:Nperc){
reassign<-round(count.healthy*(percentages[i]/100))
identification<-c(rep(F,count.healthy-
reassign),rep(T,count.infected+reassign))
vector.ident[[i]]<-identification
}
which.test<-paste(“One-tailed permutation test for contaminated data was
executed. \nProportion of differences (mean of non-infected - mean of
infected)”,
ifelse(set.higher==T,“\nhigher”,“\nlower”),
“than the observed difference is returned as an equivalent \nof p-
value.\n”,collapse=“ ”)
#Sorts healthy individuals to indicate possible false-negatives
trait2<-
c(sort(trait[infected==F],decreasing=higher.healthy),trait[infected==T])
dist.reals<-1:Nperc
names(dist.reals)<-paste(as.character(percentages), “%”)
contamination<-paste(as.character(percentages), “%”)
names(contamination)<-paste(as.character(percentages), “%”)
mean.healthy<-dist.reals
mean.infected<-dist.reals
sd.healthy<-dist.reals
sd.infected<-dist.reals
N.healthy<-dist.reals
N.infected<-dist.reals
mean.dist.perm<-dist.reals
p.vals.perm<-dist.reals
#Compute group means in non-permuted sample
for(i in 1:Nperc){
mean.healthy[i]<-mean(trait2[vector.ident[[i]]==F])
mean.infected[i]<-mean(trait2[vector.ident[[i]]==T])
sd.healthy[i]<-sd(trait2[vector.ident[[i]]==F])
sd.infected[i]<-sd(trait2[vector.ident[[i]]==T])
N.healthy[i]<-sum(vector.ident[[i]]==F)
N.infected[i]<-sum(vector.ident[[i]]==T)
dist.reals[i]<-(mean(trait2[vector.ident[[i]]==F])-
mean(trait2[vector.ident[[i]]==T]))
}
cohen<-
abs(dist.reals)/((sd.healthy*N.healthy+sd.infected*N.infected)/(N.healthy+N.
infected))
#Skewness computation
skewness.healthy<-FPskewness(trait[infected==F])
skewness.infected<-FPskewness(trait[infected==T])
skew.diff<-abs(skewness.healthy-skewness.infected)
skewness<-c(skewness.healthy,skewness.infected)
skewness<-data.frame(skewness)
names(skewness)<-“Fisher-Pearson coefficient of skewness”
rownames(skewness)<-c(“Identified as healthy”,“Identified as infected”)
signs<-sign(dist.reals)
#Permutation test with skewness add-on
perm.dist<-array(NA,dim=c(Nperc,runs))
rand.skew<-NA
for(run in 1:runs){
trait2<-sample(trait)
rand.skew[run]<-abs(FPskewness(trait2[infected==F])-
FPskewness(trait2[infected==T]))
trait2<-
c(sort(trait2[infected==F],decreasing=higher.healthy),trait2[infected==T])
for(i in 1:Nperc){
perm.dist[i,run]<-mean(trait2[vector.ident[[i]]==F])-
mean(trait2[vector.ident[[i]]==T])
}
}
skew.p<-sum(rand.skew>skew.diff)/runs
skew.higher<-ifelse(skewness.healthy>skewness.infected,“test-
negative”,“test-positive”)
skew.guess.higher<-ifelse(skewness.healthy>skewness.infected,FALSE,TRUE)
healthy.positive<-ifelse(skewness.healthy>0,TRUE,FALSE)
infected.positive<-ifelse(skewness.infected>0,TRUE,FALSE)
skew.sig<-ifelse(skew.p<0.05,TRUE,FALSE)
skew.message<-paste(
ifelse(healthy.positive==infected.positive,
paste(
“The distribution of trait value was”,
ifelse(healthy.positive,“positively”,“negatively”),
“skewed \nin both groups.”,
“The Fisher-Pearson coefficient of skewness \nwas higher
in”,skew.higher,“group.”)
,
paste(“The distribution of individuals identified as healthy \nwas skewed”,
ifelse(healthy.positive,“positively,”,“negatively,”),
“the distribution of individuals \nidentified as infected”,
ifelse(infected.positive,“positively.”,“negatively.”))
)
,
paste(“\n\nThe difference between the coefficients of skewness was”,
ifelse(skew.sig,“ \nstatistically significant.\n”,“, \nhowever,
not statistically significant.\n”),
“(Two-tailed permutation test of skewness difference \non “,runs,” runs was executed.)”,sep=“”)
,
ifelse(skew.sig==FALSE,
“\n\nThis might question the assumption of data contamination \nsince we
would expect a difference in skewness between \nthe groups in contaminated
data. \nProceed with caution.”,
paste(“\n\nThis supports the assumption of data contamination.”,
“\nBased on the difference in skewness we would assume \ncontamitation of
healthy group by false negative \nsubjects from the”,
ifelse(skew.higher==“test-positive”,“lower”,“upper”),
“tail of the distribution \nof infected individuals, which would lead to
overall”,
ifelse(skew.higher==“test-positive”,“\ndecrease”,“\nincrease”),
“of test-negative group mean.”))
,
ifelse(skew.sig==FALSE,“”,
paste(ifelse(set.higher==skew.guess.higher,
paste(“\n\nThe skewness analysis brings further support to the hypothesis
\nof”,
ifelse(set.higher,“higher”,“lower”),
“mean in non-contaminated group of healthy \nindividuals, which was used in
current permuation test \nfor contaminated data.\n”),
paste(“\n\nThe skewness analysis, however, does not support the hypothesis
\nof”,
ifelse(set.higher,“higher”,“lower”),
“mean in non-contaminated group of healthy \nindividuals, which was used in
current permuation test \nfor contaminated data. Proceed with caution.\n”)
)))
)
skewness,-rbind(skewness[1],“”,skew.p)
rownames(skewness)[c(3,4)]<-c(“”,“p-value”)
skewness[c(1,2,4),1]<-format(round(as.numeric(skewness[c(1,2,4),1]),3))
mean.dist.perm<-rowMeans(perm.dist)
if(higher.healthy==T){
for(i in 1:Nperc){
p.vals.perm[i]<-sum(dist.reals[i]<perm.dist[i,])/runs
}
}else{
for(i in 1:Nperc){
p.vals.perm[i]<-sum(dist.reals[i]>perm.dist[i,])/runs
}
}
mean.healthy<-format(round(mean.healthy,2))
mean.infected<-format(round(mean.infected,2))
sd.healthy<-format(round(sd.healthy,2))
sd.infected<-format(round(sd.infected,2))
cohen<-format(round(cohen,2))
dist.reals<-format(round(dist.reals,2))
mean.dist.perm<-format(round(mean.dist.perm,2))
p.vals.perm<-format(round(p.vals.perm,3))
res.table<-
rbind(contamination,mean.healthy,mean.infected,dist.reals,mean.dist.perm,sd
.healthy,sd.infected,N.healthy,N.infected,cohen,p.vals.perm)
res.table<-as.data.frame(res.table)
colnames(res.table)<-NULL
rownames(res.table)<-c(“contamination”,“non-infeceted
mean”,“infected mean”,“mean difference”,“expected difference”,“non-infeceted SD”,“infected
SD”,“non-infeceted N”,“infected N”,“Cohen’s d”,“p-value”)
final.message<-ifelse(all(signs>0),
“\nThe mean difference was positive in all \nhypothesised contamination
levels”,
ifelse(all(signs<0),
“\nThe mean difference was negative in all \nhypothesised contamination
levels.”,
ifelse(signs[1]>0,
“\nThe mean difference started as positive, but turned negative \nwith
growing hypothesised contamination level. \nThe results should be
interpreted with caution. \nRunning the test that assumess the opposite
relationship \nbetween group means (higher.healthy=T) or a two tailed test
\nis worth consideration.”,
“\nThe mean difference started as negative, but turned positive \nwith
growing hypothesised contamination level. \nThe results should be
interpreted with caution. \nRunning the test that assumess the opposite
relationship \nbetween group means (higher.healthy=F) or a two tailed test
\nis worth consideration.”
)))
cat(“\nSample characteristics:\n”)
print(orig.means)
cat(paste(“\n”,higher.report,“\n\n”,sep=“”))
cat(set.report1)
if(skewness.analysis==T){
cat(“\n\nSkewness report:\n”)
print(skewness)
cat(“\n”)
cat(skew.message)
}
cat(“\n\nPermutation test for contaminated data:\n”)
cat(paste(set.report2,“\n\n”,sep=“”))
cat(which.test)
cat(paste(“\n”,runs,“sample permutations were performed.\n”))
print(res.table)
cat(paste(final.message,“\n\n”,sep=“”))
results<-
list(orig.means,higher.report,set.report1,skewness,skew.message,set.report2
,which.test,res.table,final.message)
return(invisible(results))
}
###Two-tailed version of the test:
contamination_perm_test_two<-
function(trait,identification,percentages=c(0,5,10),higher.healthy=(mean(tr
ait[identification==F])>mean(trait[identification==T])),runs=10000,skewness
.analysis=F){
if(length(trait)!=length(identification)){
stop(“The vectors of trait values and infection indication are of different
lengths.”)
}
higher<-(mean(trait[identification==F])>mean(trait[identification==T]))
set.higher<-higher.healthy
orig.means<-tapply(trait,identification,mean)
orig.means<-data.frame(orig.means)
names(orig.means)<-“Original mean values”
rownames(orig.means)[which(rownames(orig.means)==“FALSE”)]<-“Identified as
healthy”
rownames(orig.means)[which(rownames(orig.means)==“TRUE”)]<-“Identified as
infected”
higher.report<-ifelse(higher==T,
“In original sample, individuals identified as healthy showed higher
\naverage trait value.”,
“In original sample, individuals identified as infected showed higher
\naverage trait value.”
)
concord<-ifelse(higher==set.higher,“Consequently,”,“Despite that,”)
set.report1<-paste(concord,ifelse(set.higher==T,
“healthy individuals were hypothesised to have higher \naverage trait value
in a contamination-free sample. \n”,
“infected individuals were hypothesised to have higher \naverage trait value
in a contamination-free sample. \n”
))
set.report2<-paste(ifelse(set.higher==T,
“\nFor each contamination level respective proportion of seronegative
\nindividuals with lowest trait value was relabeled as seropositive \nin
original sample as well as in each permutation test run.”,
“\nFor each contamination level respective proportion of seronegative
\nindividuals with highest trait value was relabeled as seropositive \nin
original sample as well as in each permutation test run.”
))
trait<-c(trait[identification==F],trait[identification==T])
infected<-sort(identification)
count.healthy<-sum(!identification)
count.infected<-sum(identification)
Nperc<-length(percentages)
vector.ident<-list()
for(i in 1:Nperc){
reassign<-round(count.healthy*(percentages[i]/100))
identification<-c(rep(F,count.healthy-
reassign),rep(T,count.infected+reassign))
vector.ident[[i]]<-identification
}
which.test<-paste(“Two-tailed permutation test for contaminated data was
executed. \nProportion of differences (in absolute value)”,
“higher than the \nobserved difference is returned as an equivalent of p-
value.\n”,collapse=“ ”)
#Sorts healthy individuals to indicate possible false-negatives
trait2<-
c(sort(trait[infected==F],decreasing=higher.healthy),trait[infected==T])
dist.reals<-1:Nperc
names(dist.reals)<-paste(as.character(percentages), “%”)
contamination<-paste(as.character(percentages), “%”)
names(contamination)<-paste(as.character(percentages), “%”)
mean.healthy<-dist.reals
mean.infected<-dist.reals
sd.healthy<-dist.reals
sd.infected<-dist.reals
N.healthy<-dist.reals
N.infected<-dist.reals
mean.dist.perm<-dist.reals
p.vals.perm<-dist.reals
#Compute group means in non-permuted sample
for(i in 1:Nperc){
mean.healthy[i]<-mean(trait2[vector.ident[[i]]==F])
mean.infected[i]<-mean(trait2[vector.ident[[i]]==T])
sd.healthy[i]<-sd(trait2[vector.ident[[i]]==F])
sd.infected[i]<-sd(trait2[vector.ident[[i]]==T])
N.healthy[i]<-sum(vector.ident[[i]]==F)
N.infected[i]<-sum(vector.ident[[i]]==T)
dist.reals[i]<-(mean(trait2[vector.ident[[i]]==F])-
mean(trait2[vector.ident[[i]]==T]))
}
cohen<-
abs(dist.reals)/((sd.healthy*N.healthy+sd.infected*N.infected)/(N.healthy+N
.infected))
#Skewness computation
skewness.healthy<-FPskewness(trait[infected==F])
skewness.infected<-FPskewness(trait[infected==T])
skew.diff<-abs(skewness.healthy-skewness.infected)
skewness<-c(skewness.healthy,skewness.infected)
skewness<-data.frame(skewness)
names(skewness)<-“Fisher-Pearson coefficient of skewness”
rownames(skewness)<-c(“Identified as healthy”,“Identified as infected”)
signs<-sign(dist.reals)
#Permutation test with skewness add-on
perm.dist<-array(NA,dim=c(Nperc,runs))
rand.skew<-NA
for(run in 1:runs){
trait2<-sample(trait)
rand.skew[run]<-abs(FPskewness(trait2[infected==F])-
FPskewness(trait2[infected==T]))
trait2<-
c(sort(trait2[infected==F],decreasing=higher.healthy),trait2[infected==T])
for(i in 1:Nperc){
perm.dist[i,run]<-mean(trait2[vector.ident[[i]]==F])-
mean(trait2[vector.ident[[i]]==T])
}
}
skew.p<-sum(rand.skew>skew.diff)/runs
skew.higher<-ifelse(skewness.healthy>skewness.infected,“test-
negative”,“test-positive”)
skew.guess.higher<-ifelse(skewness.healthy>skewness.infected,FALSE,TRUE)
healthy.positive<-ifelse(skewness.healthy>0,TRUE,FALSE)
infected.positive<-ifelse(skewness.infected>0,TRUE,FALSE)
skew.sig<-ifelse(skew.p<0.05,TRUE,FALSE)
skew.message<-paste(
ifelse(healthy.positive==infected.positive,
paste(
“The distribution of trait value was”,
ifelse(healthy.positive,“positively”,“negatively”),
“skewed \nin both groups.”,
“The Fisher-Pearson coefficient of skewness \nwas higher
in”,skew.higher,“group.”)
,
paste(“The distribution of individuals identified as healthy \nwas skewed”,
ifelse(healthy.positive,“positively,”,“negatively,”),
“the distribution of individuals \nidentified as infected”,
ifelse(infected.positive,“positively.”,“negatively.”))
)
,
paste(“\n\nThe difference between the coefficients of skewness was”,
ifelse(skew.sig,“ \nstatistically significant.\n”,“, \nhowever, not
statistically significant.\n”),
“(Two-tailed permutation test of skewness difference \non ”,runs,“ runs was
executed.)”,sep=“”)
,
ifelse(skew.sig==FALSE,
“\n\nThis might question the assumption of data contamination \nsince we
would expect a difference in skewness between \nthe groups in contaminated
data. \nProceed with caution.”,
paste(“\n\nThis supports the assumption of data contamination.”,
“\nBased on the difference in skewness we would assume \ncontamitation of
healthy group by false negative \nsubjects from the”,
ifelse(skew.higher==“test-positive”,“lower”,“upper”),
“tail of the distribution \nof infected individuals, which would lead to
overall”,
ifelse(skew.higher==“test-positive”,“\ndecrease”,“\nincrease”),
“of test-negative group mean.”))
,
ifelse(skew.sig==FALSE,“”,
paste(ifelse(set.higher==skew.guess.higher,
paste(“\n\nThe skewness analysis brings further support to the hypothesis
\nof”,
ifelse(set.higher,“higher”,“lower”),
“mean in non-contaminated group of healthy \nindividuals, which was used in
current permuation test \nfor contaminated data.\n”),
paste(“\n\nThe skewness analysis, however, does not support the hypothesis
\nof”,
ifelse(set.higher,“higher”,“lower”),
“mean in non-contaminated group of healthy \nindividuals, which was used in
current permuation test \nfor contaminated data. Proceed with caution.\n”)
)))
)
skewness<-rbind(skewness[1],“”,skew.p)
rownames(skewness)[c(3,4)]<-c(“”,“p-value”)
skewness[c(1,2,4),1]<-format(round(as.numeric(skewness[c(1,2,4),1]),3))
mean.dist.perm<-rowMeans(perm.dist)
if(higher.healthy==T){ for(i in 1:Nperc){
p.vals.perm[i]<-sum(dist.reals[i]<perm.dist[i,])/runs
}
}else{
for(i in 1:Nperc){
p.vals.perm[i]<-sum(dist.reals[i]>perm.dist[i,])/runs
}
}
mean.healthy<-format(round(mean.healthy,2))
mean.infected<-format(round(mean.infected,2))
sd.healthy<-format(round(sd.healthy,2))
sd.infected<-format(round(sd.infected,2))
cohen<-format(round(cohen,2))
dist.reals<-format(round(dist.reals,2))
mean.dist.perm<-format(round(mean.dist.perm,2))
p.vals.perm<-format(round(p.vals.perm,3))
res.table<-
rbind(contamination,mean.healthy,mean.infected,dist.reals,mean.dist.perm,sd
.healthy,sd.infected,N.healthy,N.infected,cohen,p.vals.perm)
res.table<-as.data.frame(res.table)
colnames(res.table)<-NULL
rownames(res.table)<-c(“contamination”,“non-infeceted mean”,“infected
mean”,“mean difference”,“expected difference”,“non-infeceted SD”,“infected
SD”,“non-infeceted N”,“infected N”,“Cohen’s d”,“p-value”)
final.message<-paste(“\nTwo-tailed permutation test for contaminated data
was executed. \nIt was assumed that”,
ifelse(set.higher==T,“healthy”,“infected”),
“individuals have on average higher \ntrait value if all false negative
individuals are relocated correctly.”)
cat(“\nSample characteristics:\n”)
print(orig.means)
cat(paste(“\n”,higher.report,“\n\n”,sep=“”))
cat(set.report1)
if(skewness.analysis==T){
cat(“\n\nSkewness report:\n”)
print(skewness)
cat(“\n”)
cat(skew.message)
}
cat(“\n\nPermutation test for contaminated data:\n”)
cat(paste(set.report2,“\n\n”,sep=“”))
cat(which.test)
cat(paste(“\n”,runs,“sample permutations were performed.\n”))
print(res.table)
cat(paste(final.message,“\n\n”,sep=“”))
results<-
list(orig.means,higher.report,set.report1,skewness,skew.message,set.report2
,which.test,res.table,final.message)
return(invisible(results))
}
~~~

### Supplement 2

#### Skewness analysis

~~~
#####################################################
####hit ctrl+a and ctrl+r to install the function####
#####################################################
#This script contains skewness_comparison() function
#skewness_comparison(trait,identification,percentages=seq(0,50,1),higher.healthy=(mean(trait[identification==F])>mean(trait[identification==T])),runs= 10000)
#Arguments are described below.
#It is necessary to define funtion calculating skewness index first:
##Fuction that returns Fisher-Pearson coefficient of skewness.
##Input is a vector of numerical values.
FPskewness<-function(x){
return((sum((x-mean(x))^3)/length(x))/((sqrt(sum((x-
mean(x))^2)/length(x)))^3))
}
###Skewness comparison
###Function that reports how relocation of seronegative individuals
changes the skewnees of the distribution in both seronegtive and seropositive
groups.
###trait - Numerical vector of trait values
###identification - Logical vector of assumed presence (T) or absence (F)
of infection
###percentages - Numerical vector of percentages of false negative amongst
negative subjects (contamination levels) for which the permutation test for
contaminated data will be run
###higher healthy - Logical. Indicates whether we assume the healthy
individuals to show higher (T) or lower (F) trait values. When not
specified, the script assumes this relationship based of group means with
no hypothesised contamination.
######It allows us to use the difference between the groups (not in
absolute values) in permutation test. In this scenario the seropositive
group mean is substracted from seronegative group mean and the one-tailed
permutation test is conducted accordingly.
###runs - Number of resamplings used in permutation test
skewness_comparison<-
function(trait,identification,percentages=seq(0,50,1),higher.healthy=(mean(
trait[identification==F])>mean(trait[identification==T])),runs=10000){
if(length(trait)!=length(identification)){
stop(“The vectors of trait values and infection indication are of different
lengths.”)
}
higher<-(mean(trait[identification==F])>mean(trait[identification==T]))
set.higher<-higher.healthy
orig.means<-tapply(trait,identification,mean)
orig.means<-data.frame(orig.means)
names(orig.means)<-“Original mean values”
rownames(orig.means)[which(rownames(orig.means)==“FALSE”)]<-“Identified as healthy”
rownames(orig.means)[which(rownames(orig.means)==“TRUE”)]<-“Identified as infected”
higher.report<-ifelse(higher==T,
“In original sample, individuals identified as healthy showed higher
\naverage trait value.”,
“In original sample, individuals identified as infected showed higher
\naverage trait value.”
)
concord<-ifelse(higher==set.higher,“Consequently,”,“Despite that,”)
set.report1<-paste(concord,ifelse(set.higher==T,
“healthy individuals were hypothesised to have higher \naverage trait value
in a contamination-free sample. \n”,
“infected individuals were hypothesised to have higher \naverage trait
value in a contamination-free sample. \n”
))
run.report<-paste(“Two-tailed permutation test of skewness difference \non
”,runs,“ runs was executed on each contamination level.\n\n”,sep=“”)
set.report2<-paste(“\nFor each contamination level respective proportion of
seronegative \nindividuals with”,
ifelse(set.higher==T,“lowest”,“highest”),
“trait value was relabeled as seropositive \nand the difference between the
group skewness was measured. \nReferential skewness differences from
permutation runs were based \non random non-contaminated sample with group
sizes corresponding \nto respective contamination levels.\n\n”
)
trait<-c(trait[identification==F],trait[identification==T])
infected<-sort(identification)
count.healthy<-sum(!identification)
count.infected<-sum(identification)
Nperc<-length(percentages)
vector.ident<-list()
for(i in 1:Nperc){
reassign<-round(count.healthy*(percentages[i]/100))
identification<-c(rep(F,count.healthy-
reassign),rep(T,count.infected+reassign))
vector.ident[[i]]<-identification
}
#Sorts healthy individuals to indicate possible false-negatives
trait2<-
c(sort(trait[infected==F],decreasing=higher.healthy),trait[infected==T])
skewness.healthy<−1:Nperc
names(skewness.healthy)<-paste(as.character(percentages), “%”)
skewness.infected<-skewness.healthy
contamination<-paste(as.character(percentages), “%”)
names(contamination)<-paste(as.character(percentages), “%”)
#Compute group means in non-permuted sample
for(i in 1:Nperc){
skewness.healthy[i]<-FPskewness(trait2[vector.ident[[i]]==F])
skewness.infected[i]<-FPskewness(trait2[vector.ident[[i]]==T])
}
skew.diff<-abs(skewness.healthy-skewness.infected)
#Permutation test of skewness difference
skew.diff.perm<-array(NA,dim=c(Nperc,runs))
for(run in 1:runs){
trait2<-sample(trait)
for(i in 1:Nperc){
skew.diff.perm[i,run]<-abs(FPskewness(trait2[vector.ident[[i]]==F])-
FPskewness(trait2[vector.ident[[i]]==T]))
}
}
p.vals.skew<-NA
for(i in 1:Nperc){
p.vals.skew[i]<-sum(skew.diff[i]<skew.diff.perm[i,])/runs
}
guess.perc<-percentages[which(skew.diff==min(skew.diff))]
possible<-percentages[p.vals.skew>0.05]
if(length(possible)==0){
ok.report<-paste(“\nThe difference in Fisher-Pearson coefficients of
skewness between \ngroups was significant at all investigated contamination
levels.\nThis may be caused by extreme proportion of false negative
\nindividuals, insufficient number of relocated fractions, \ndifferent
shapes of distributions of healthy and infeceted \nindividuals, or, most
likely, by wrong setting of higher group \nmean in non-contaminated sample.
\nTry running this comparison with parameter higher.healthy
=”,ifelse(higher.healthy==TRUE,“FALSE”,“TRUE”),“\n\n”)
}else{
if(min(possible)==max(possible)){
ok.report<-paste(“\nThe difference in Fisher-Pearson coefficients of
skewness between \ngroups was not significant at”,
min(possible),
“% of relocated individuals.\n\n”)
}else{
ok.report<-paste(“\nThe difference in Fisher-Pearson coefficients of
skewness between \ngroups was not significant between”,
min(possible),
“% and”,
max(possible),
“% of relocated individuals.\n\n”)
}
}
best.guess<-paste(“The difference between group skewness was smallest
\nwhen”,
guess.perc,
“% of seronegative individuals with”,
ifelse(higher.healthy,“lowest”,“highest”),
“trait value \nwas relocated to seropositive group.\n\n”)
skewness.healthy<-format(round(skewness.healthy,2))
skewness.infected<-format(round(skewness.infected,2))
p.vals.skew<-format(round(p.vals.skew,3))
skewness.res<-
rbind(contamination,skewness.healthy,skewness.infected,“”,p.vals.skew)
skewness.res<-as.data.frame(skewness.res)
colnames(skewness.res)<-NULL
rownames(skewness.res)<-c(“contamination”,“skewness healthy”,“skewness
infected”,“”,“p-value”)
cat(“\nSample characteristics:\n”)
print(orig.means)
cat(paste(“\n”,higher.report,“\n\n”,sep=“”))
cat(set.report1)
cat(set.report2)
cat(run.report)
cat(paste(“Skewness comparison:”,“\n”,sep=“”))
print(skewness.res)
cat(ok.report)
cat(best.guess)
results<-list(orig.means,higher.report,set.report1,skewness.res)
return(invisible(results))
}
~~~

### Supplement 3

#### Example

~~~
###################
###Exemplar runs###
###################
#Both functions contamination_perm_test() and skewness_comparison() must be
installed, run respective scripts
#You can generate data with known proportion of false negative individuals
with this script and try permutation test for contaminated data on them.
count<-1000 #sets sample size
inf.prop<-0.5 #sets proportion of seropositive individuals
fixed.effect<-(−3) #sets the effect of infection on simulated trait
healthy.average<-101.5 #sets the average trait value in noninfected group
sd<-12 #sets the standard deviation of within group
false.negatives<-5 #sets proportion of false negative individuals
###Computes counts in respektive groups using set properties
count.infected<-round(inf.prop*count)
count.healthy<-count-count.infected
###Creates a variables that indicates infection
infected<-c(rep(F,count.healthy),rep(T,count.infected))
#Calculates how many false negative individuals will be in the seronegative
group
reassign<-round(count.healthy*(false.negatives/100))
#Creates a vector of actual iinfection
really.infected<-c(rep(F,count.healthy-
reassign),rep(T,count.infected+reassign))
#Generates the population (all healthy individuals)
trait<-rnorm(count,healthy.average,sd)
#modifies really infected individuals
trait[(sum(!really.infected)+1):count]<-
trait[(sum(!really.infected)+1):count]+fixed.effect
#sorts really infeceted individuals such that most changed ones are close
in the vector to healthy ones i.e. are marked as test-negative
trait<- c(trait[really.infected==F],sort(trait[really.infected==T],decreasing=(sign
(fixed.effect)==1)))
#scrambles the vectors, along the same random vector
scramble<-sample(1:count)
infected<-infected[scramble]
trait<-trait[scramble]
##########################
###Trial data are ready###
##########################
#Executes permutation test for contaminated data with default argument
values
contamination_perm_test(trait,infected)
#Executes permutation test for contaminated data and skewness analysis
contamination_perm_test(trait,infected,skewness.analysis=T)
#Executes permutation test for contaminated data, hypothesised diffrence is
specified by hand. This comes useful when you have a reason to suspect
paradoxical switch in group means.
contamination_perm_test(trait,infected,higher.healthy=F)
#Executes permutation test on only 100 permutation runs per test - it is
quicker, but less accurate
contamination_perm_test(trait,infected,runs=100)
#Executes two tailed permutation test for contaminated dat
contamination_perm_test(trait,infected,two.tailed=T)
#Executes skewness comparison to estimate proportion and distribution tail
of possible contamination prior to the test
skewness_comparison(trait,infected)
#Executes permutation test for contaminated data, levels of contamination
are specified by hand
contamination_perm_test(trait,infected,percentages=seq(1,15,1))
~~~

